# Standing balance in aging is robust against head rotation during visual tracking

**DOI:** 10.1101/2022.07.30.502138

**Authors:** Petros Georgiadis, Konstantinos Chatzinikolaou, Dimitrios Voudouris, Jaap Van Dieen, Vassilia Hatzitaki

## Abstract

Standing balance is relatively more unstable when visually pursuing a moving target than when fixating a stationary one. These effects are common across age groups if the head is restrained during visual task performance. The present study focused on the role of the head motion on standing balance during the pursuit of a moving target as a function of age. Three predictions were tested: a) standing balance is compromised to a greater extent in older than young adults by gaze target pursuit compared to fixation, b) older adults pursue a moving target with greater and more variable head rotation than young adults, and c) greater and more variable head rotation during the gaze pursuit task is associated with greater postural sway. Twenty-two (22) older (age: 71.7±8.1, 12 M / 10 F) and twenty-three (23) young adults (age: 23.6±2.5, 12 M / 11 F) stood on a force plate in front of a 60-inch monitor while performing two visual tasks: fixation at a stationary target and gaze pursuit of a horizontally moving target. Centre of pressure (CoP) and head kinematics were synchronously recorded with the Vicon motion analysis system, while head-unconstrained gaze was captured by the Pupil Labs Invisible mobile tracking system. Postural sway, reflected in the interquartile CoP range and the root mean square (RMS) of CoP velocity increased during the gaze pursuit compared to the fixation task (p<.05), and this effect was more pronounced in older than young participants (p<.05). Older adults pursued the moving target employing more variable (p=.022) head yaw rotation than young participants although the amplitude of head rotation was not systematically different between groups (p=. 077). The amplitude and variance of head yaw rotation did not correlate with postural sway measures. Results suggest that older adults may engage more variable head rotation when tracking a moving target to compensate for age-related deficits in eye smooth pursuit movement. However, this strategy does not seem to compromise standing balance.

## Introduction

Balance control is crucial for the maintenance of autonomous function and mobility in old age. Older adults show increased reliance on visual input for balance control to compensate for their age-related decline in muscle proprioception and somatosensation (Eikema et al. 2013). At the same time, the majority of falls in older adults occurs during a postural transition (sit to stand, stand to walk transitions) or weight transfer (Robinovitch et al. 2010), actions that require rapid reorientation of the gaze from the current to the next spatial location. When re-orienting gaze while standing or walking on a treadmill, older adults exhibit greater head rotation compared to young adults (Cinelli et al. 2008). This is accompanied by concurrent shoulder and hip rotations to the intended direction of motion pointing to an en-bloc balance strategy, which could compromise stability. Moreover, in real world conditions such as walking down a hallway with free gaze or fixating an earth-fixed object while walking, older adults employ fewer, smaller, and slower saccadic eye movements (Dowiasch et al. 2015) which could negatively affect visual perception and early motion processing required for successful balance control. These age-related gaze alterations result in less precise and more variable steps during locomotion increasing the risk of falling (Chapman & Hollands 2006).

Tracking of a smoothly and slowly moving target (<100°/s) could also compromise balance. Performance of smooth pursuit eye movements increases postural sway relative to target fixation (Guerraz & Bronstein 2008). However, the destabilizing effect does not seem to be age-specific, because young and older adults increased mediolateral center of pressure (CoP) sway to a similar extent when tracking a target moving slowly on a stationary background (Thomas et al. 2016). Similarly, walking into a room while fixating a stationary person or tracking a moving one, decreased balance control, but again, this effect was similar in young and older females (Thomas et al. 2018). The absence of an age-specific effect of smooth pursuit on postural sway could be due to the fact that most eye tracking systems require the head to remain stationary during visual task performance (Glasauer et al. 2005; Stoffregen 2006; Thomas et al. 2016; Thomas et al. 2018). In the real world, although an object can be visually tracked by moving the eyes with or without concurrent head motion, accurate vision is mostly achieved through the coordinated movement of the eyes and the head (Kowler 2011). Head rotation and eye movements can occur simultaneously to produce larger gaze excursions extending the initial visual field restrictions (Freedman & Sparks 1997). This can have important implications on postural sway, as older adults seem to engage their head more strongly when shifting gaze (Cinelli et al., 2008), which might be related to their reduced amplitude and gain of smooth pursuit eye movements (Paige 1994; Zackon & Sharpe 2009; Maruta et al. 2017). A rising concern however is whether this possible extra head motion challenges older adults’ standing balance, increasing the risk of falling in daily activities that require visual tracking. Alternatively, it does not seem unreasonable to hypothesize that older adults exploit the extra head rotation to gain additional sensory input from head-neck proprioception and vestibular signals to facilitate postural control. Yet, it is not known how pursuing a moving target impacts older adults’ standing balance when the head is free to move. In the literature, we found only one study investigating age effects on smooth gaze pursuit when the head is free during standing (Paquette & Fung 2011), which showed an age-related decline in tracking accuracy and gaze-target coupling, but did not report any effects on standing balance.

The present study was designed to address the above issues focusing on the following aims: a) investigate the effect of gaze fixation and smooth pursuit on older and young adults’ standing balance, b) identify gaze pursuit strategies of young and older adults based on the amplitude and variance of head yaw rotation, and c) assess the relationship between head motion and standing balance. Static balance was assessed during gaze fixation on a stationary target and during gaze pursuit of a target moving horizontally on a stationary (black) background. Three predictions were made: a) standing balance is compromised to a greater extent in older than young adults by the gaze pursuit when compared to the gaze fixation task, b) older adults pursue the moving target with greater head yaw rotation, and c) the greater age-induced instability is related to greater head yaw rotation for pursuing the moving target.

## Materials and methods

### Participants

Twenty-two (22) older (age: 71.7±8.1, 12 M / 10 F) and twenty-three (23) young (age: 23.6±2.5, 12 M / 11 F) adults participated in the study. Young participants were volunteers among university students, whereas the older participants were recruited from day care community centers for older adults. Inclusion criteria were: (1) no symptoms suggestive of eye disorder that could affect the reliability of the gaze measurement; (2) no orthopedic diagnoses preventing standing; (3) no medications known to affect the nervous system or balance. Participants were assessed with the Mini Mental State Examination and for visual acuity using a Snellen board. Younger and older participants achieved a score ≥ 24 and ≥ 20/100 respectively, both considered acceptable for inclusion in the study. Participants were informed about the purpose and the experimental protocol of the study and gave written informed consent. Experimental procedures were approved by Aristotle University of Thessaloniki Ethics Committee on Human Research in accordance with the Declaration of Helsinki (Approval number 100/2022).

### Apparatus, visual stimuli, and task protocol

A force platform (Balance Plate 6501, Bertec, USA, sampling rate: 100Hz) recorded the vertical ground reaction force and moments in the anteroposterior and mediolateral direction to calculate the Center of Pressure displacement (CoP). A TV screen (LG 60LA620S-ZA, 60 inches, 1.5 m horizontal x 0.8 m vertical; screen refresh rate: 60Hz) was positioned 1.95 m in front of the participant centered at eye level. Kinematic data were captured using a 10-camera system (Vicon Motion Systems, Oxford, UK, sampling rate: 100 Hz). Reflective markers were attached at the following anatomical landmarks: a) head (two markers on the forehead) b) right and left acromion processes, c) right and left trochanter. The Vicon’s Software Development Kit (SDK) was used to establish communication between the Vicon system software (Nexus v. 1.8.5) and LabView (version 8.6, National Instruments Corporation, 2008), allowing the visual stimuli presentation and synchronous sampling and digitization of the force, marker kinematics and target motion signals via Vicon’s data acquisition board (MX Giganet). The Pupil Invisible mobile eye tracking system (Pupil Labs GmbH; sampling rate: 200 Hz) was used to capture the gaze parameters. The synchronization of the Vicon motion analysis and the visual tracking systems was based on the synchronous recording of a sound that was generated by the LabView custom software developed to control the experiment. The sound was stored as an analog signal in both Nexus and the eye tracking software and used to synchronize the time series offline.

The visual stimulus was a red circular (disk shaped) target (diameter equivalent to 1.46° of visual angle) projected onto a stationary totally black background, which occupied the full screen dimensions (Fig1). The vertical center of the TV screen was adjusted to the participant’s eye level. The target was always presented 10% below the vertical center of the screen to correspond with the mean standard line of sight and could be stationary (fixation task) or move horizontally covering 90% of the screen’s width (visual angle of 31.9° or ± 15.95° from the midline) at a constant velocity of 15.96°/s (tracking task). Gaze shifts of <15° are commonly achieved without rotation of the head (Hallet, 1986; Stoffregen et al 2006). The target’s motion signal was a sine wave created in LabView (version 8.6) of a frequency of 0.25 Hz which gave 15 cycles of horizontal motion in 60 s.

**Fig. 1:**
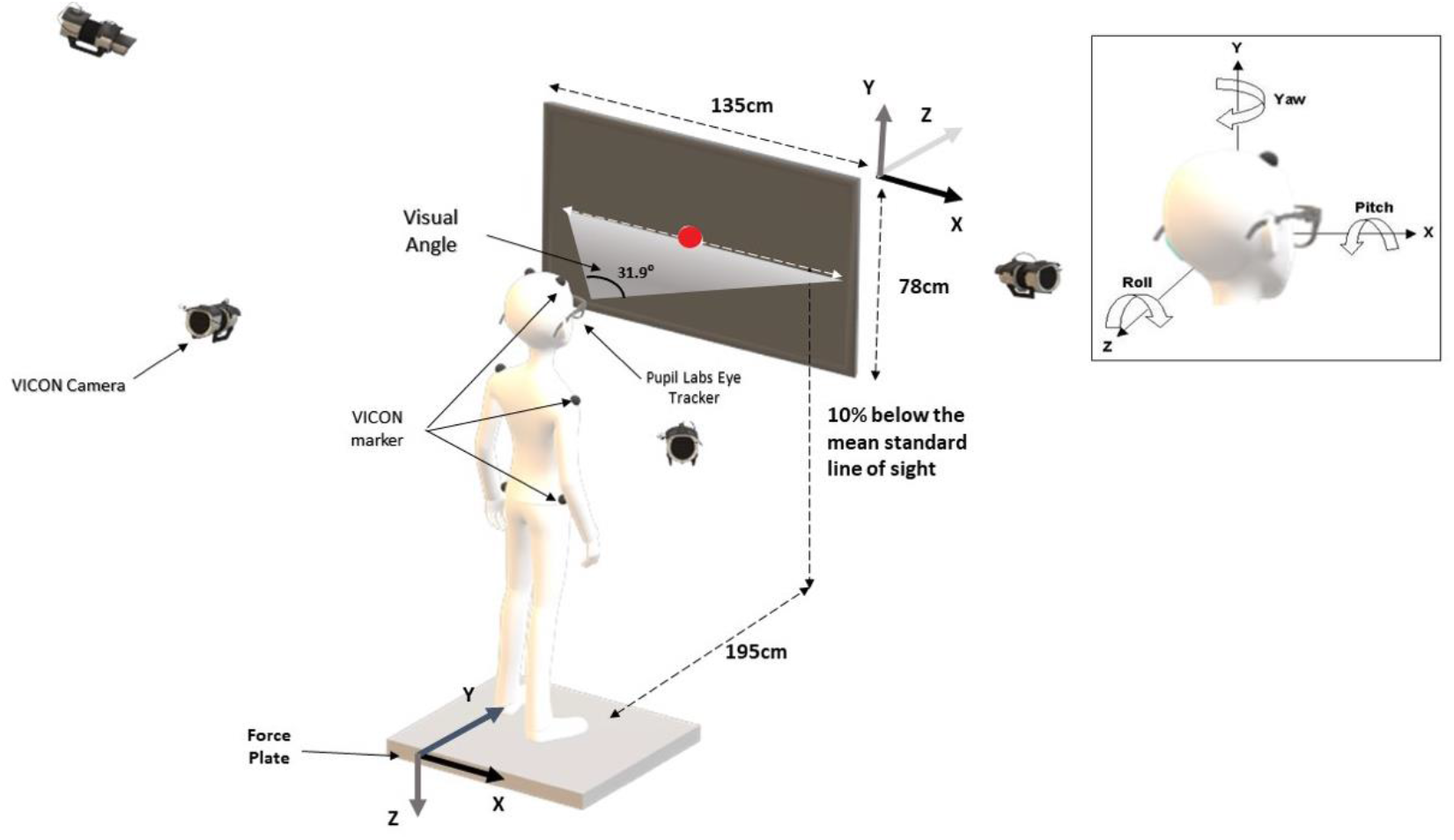
Experimental setup and task. The participant stood on a force platform in front of a TV screen and fixated a red circular target displayed on a black background (30s) or followed with gaze the horizontal motion of the target (60s). Ground reaction forces, gaze and head kinematics were synchronously recorded.

Participants stood in a bipedal upright position at the center of the force platform, barefooted with feet parallel, keeping intermalleolar distance to 10% of the shoulder-to-shoulder distance and with arms free hanging on body sides (Fig 1). While standing, they were asked to perform one of the following visual tasks in a randomized order: a) fixate the stationary target for 30 s (fixation task), b) pursue the horizontally moving target for 60 s (gaze pursuit task). No instruction about head stabilization was given. Instead, participants were free to move body segments at their convenience to accurately fixate on or pursue the visual target. The two conditions were part of a larger study protocol.

### Data analysis

Force, kinematic and gaze timeseries were processed in MATLAB (2019b) using custom-made scripts. For the data analysis, the first and last 5 seconds of each (fixation and pursuit) trial were disregarded to avoid signal irregularities due to transient effects of task familiarization and fatigue.

### CoP

Medial/lateral (ML) and anterior/posterior (AP) CoP timeseries were first computed from the vertical ground reaction force vector and the two moments. The CoP time series were then filtered with a 4^th^ order low-pass Butterworth filter (cut-off frequency 5Hz) and differentiated to calculate CoP velocity. Postural sway was summarized with the following dependent variables: i) interquartile CoP range in the AP and ML direction (CoP range) and ii) Root Mean Square of the CoP velocity (RMS CoPvel) in the AP and ML direction.

### Head Kinematics

For the calculation of head rotation in the yaw plane, the Cartesian 2D linear position coordinates (x,y) of the right and left head markers were smoothened with a digital low pass Butterworth filter (cut-off 5Hz) and then converted to angular coordinates using the arc tangent function (Winter, 1999). For the fixation task, the head range of motion (angular rotation) around the yaw axis was calculated by subtracting the minimum from the maximum value of the head angular displacement time series. For the tracking task, the head range of motion was calculated for each target cycle and then this value was averaged across the 13 cycles of target motion (after subtracting the first and last cycle of the trial) to get a) the mean and b) the standard deviation (SD) of the head rotation amplitude as a measure of the amplitude and variance of head rotation respectively.

### Gaze

To calculate the two-dimensional linear coordinates of gaze position on the monitor, Apriltag markers were placed at the 4 corners of the TV screen. Next, using an algorithm for recognizing their position, the active registration surface was calibrated (White et al. 2007). The calculation of the points on screen where the 2-eyes line of sights converged, was made through Pupil Labs Player software. The initial recording of the gaze position on the TV surface was calibrated to pixels with sampling frequency of 200Hz before it was converted into visual angle in degrees. For the visual tracking task, the spatial coupling between gaze and the target motion was assessed as the peak of the cross-correlation function (CCF) between the gaze and the target motion timeseries. A value of 1 represents perfect spatial coupling between the time series.

### Sample size estimation and statistical analysis

Sample size was estimated based on the age induced difference in RMS of CoP velocity in the ML direction during the gaze pursuit task. After collecting pilot data on 5 participants of each group in this task condition, we considered a mean expected age group difference of 0.3336 cm ± .3231 (SD) with α=0.05 and power=0.90. Using this information, we calculated that a minimum of 21 participants per group (effect size = 1.032) would be sufficient to test our hypothesis about the effect of age.

Prior to statistical testing all variable distributions were checked for normality (Kolomogorov-Smirnov test) and homogeneity of variance (Levene’s test) across groups. To test the 1^st^ hypothesis of the study, the effect of age group and visual task (fixation vs. tracking) on the postural sway variables (CoP range and RMS of CoP velocity) was investigated employing a 2 × 2 ANOVA with repeated measures on task. Estimates of effect size on all dependent measures are reported using partial η^2^. Task by group interactions were tested by pairwise comparisons between the visual tasks performed separately for each group. To test the 2^nd^ hypothesis of the study associated with age related differences in the gaze pursuit strategy, the mean and SD of head yaw rotation amplitude and the peak of the gaze-target CCF during the gaze pursuing task were compared across age groups employing t-test or non-parametric Mann-Whitney U test for independent samples. To test the 3^rd^ hypothesis of the study, Pearson’s correlations were calculated between the postural sway variables and the head yaw rotation variables during the gaze pursuit task separately for the young and old group adults.

## Results

### Fixation vs gaze pursuit effects on standing balance

All postural variables were normally (Kolomogorov-Smirnov test: p>.05) and homogeneously (Levene’s test p>.05) distributed. Fig 2 shows exemplar ML-AP CoP traces of one older and one young participant plotted for the two visual tasks.

**Fig 2:**
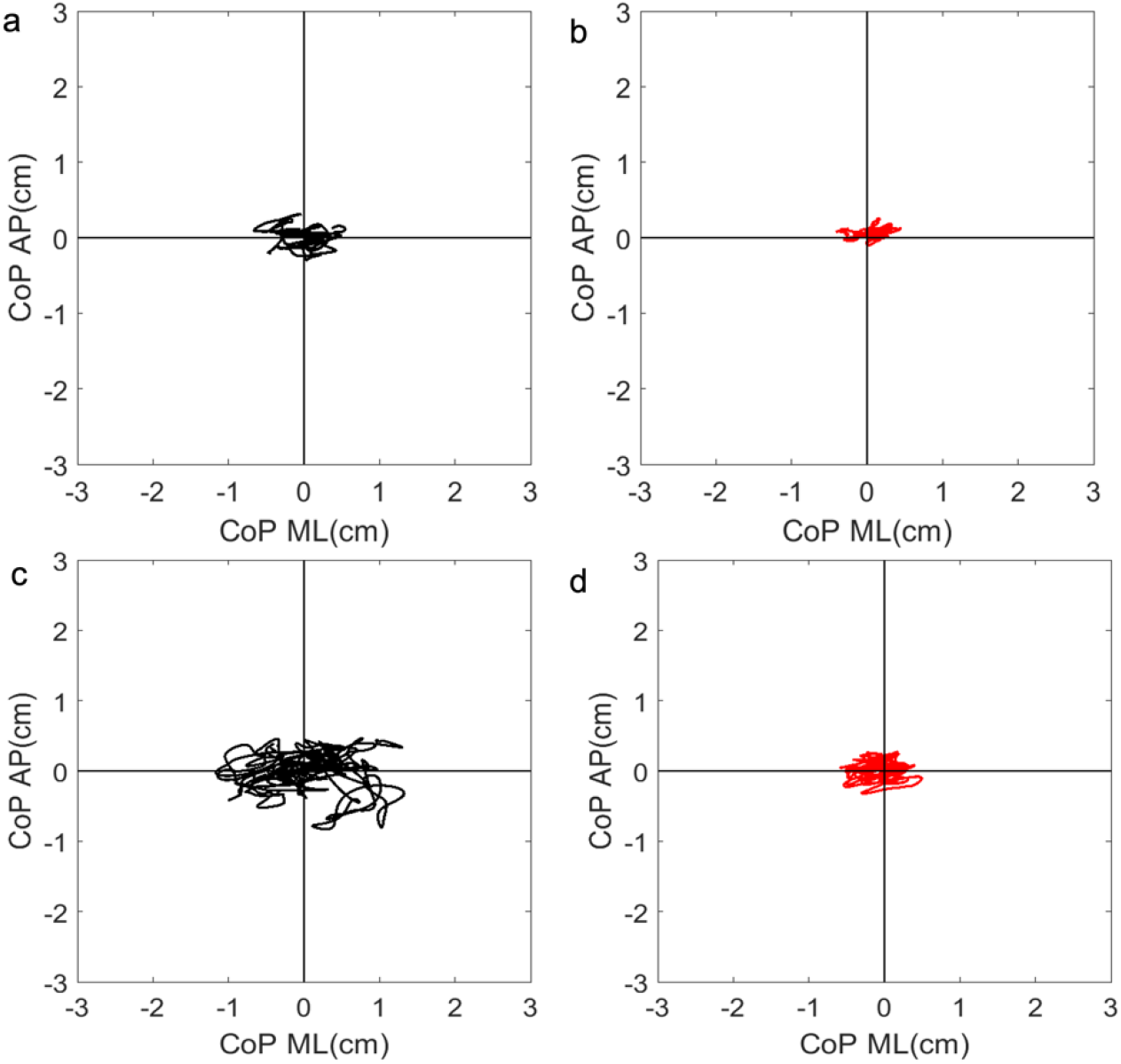
Representative CoP time series plotted in the ML vs the AP direction for the gaze fixation (a,b)) and pursuit tasks (c,d) of one old (a,c in black) and one young (d,.d in red) participant

Pursuing the moving target destabilized quiet standing when compared to gaze fixation as confirmed by a significant main effect of the task on all postural variables [CoP ML range: F(1,43)=6.85, p=.012, η2=.147, Fig 3a; CoP AP range: F(1,43)=14.29, p<.001, η2=.250, Fig. 3b; CoP RMS vel ML: F(1,43)=18.56, p<.001, η2=.301, Fig 3c; CoP RMS vel AP: F(1,43)=16.09, p<.001, η2=.272, Fig. 3d]. A significant group effect of CoP RMS vel ML [F (1,43) =18.68, p<.001, η2=.303] revealed that older adults had significantly greater variance of CoP velocity in the direction of the target’s motion. More importantly, the effect of the gaze pursuit task on CoP range was dependent on the age group as revealed by a significant age group by task interaction [CoP ML range: F (1,43) =3.96, p=.05, η2=.084; CoP AP range: F (1,43) =5.036, p=.03, η2=.105]]. Subsequent post hoc pairwise comparisons confirmed that, in older adults, CoP interquartile range was greater when pursuing the target compared to fixating to it [CoP ML range (x): (t (21) = 2.75, p=.012; CoP AP range (y): (t (21) = 3.56, p=.002], but this was not the case for young adults (all t < 1.945, all p>.05).

**Fig. 3:**
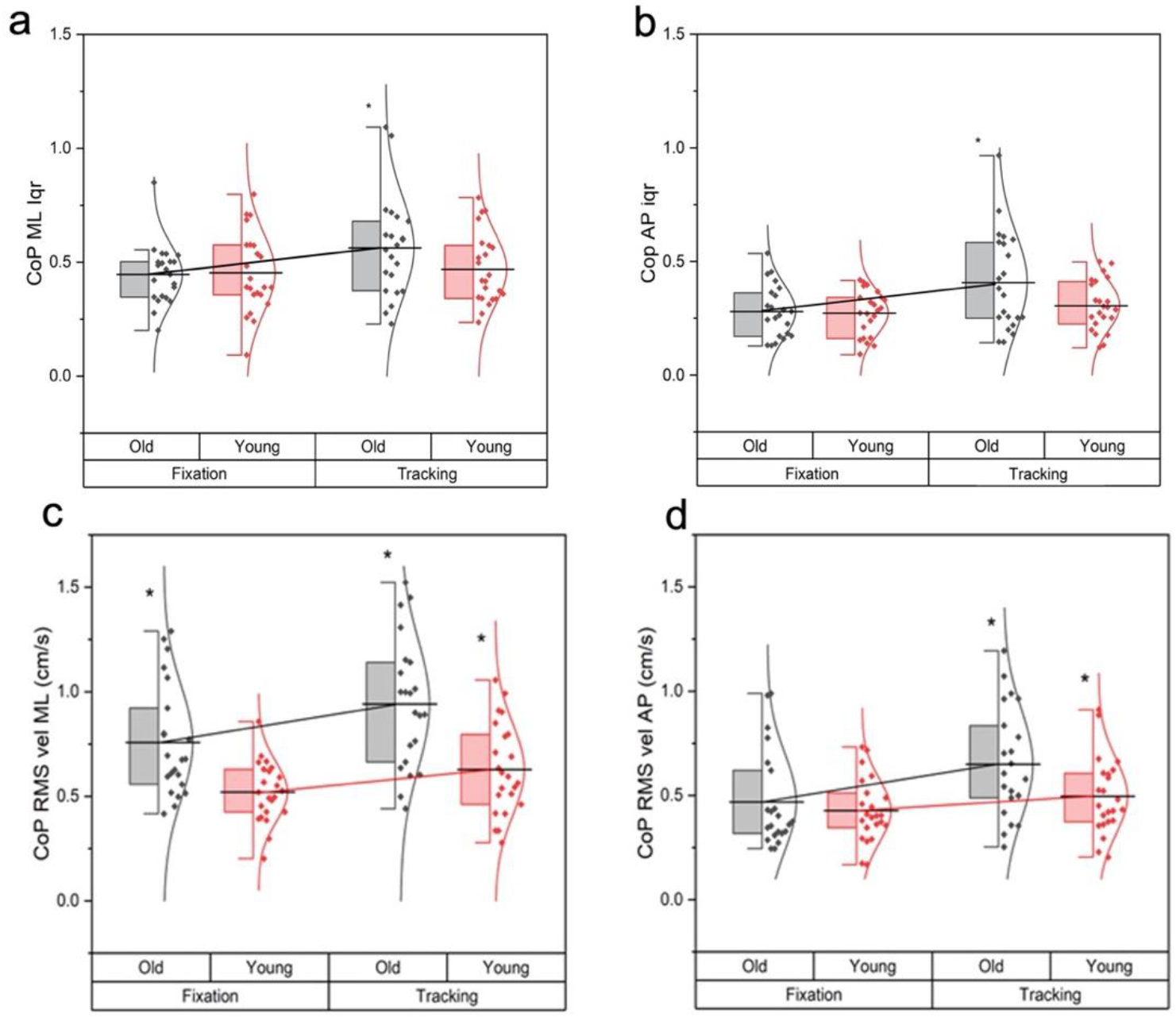
Interquartile CoP range (CoP iqr, cm)) in the mediolateral (a) and anteroposterior direction (b), RMS of CoP velocity (c,d) in the respective directions for Fixation vs Pursuit (group means + se). *: > in tracking (p<.05).

### Age-related differences in head rotation

The range of the head yaw rotation during the fixation task was <1° for all group participants regardless of age which suggests that the head was stationary during the fixation task.

Fig 4a and 4b show the individual head yaw rotation time series for the old and young group participants respectively plotted together with the group mean (thick black line). Individual values of the mean head rotation amplitude (Fig 4c) were widely spread, positively skewed (skewness old group: 1.267, young group: 1.617), not normally (Kolomogorov-Smirnov test: young group: p = .002, old group: p = .038) and non-homogeneously distributed (Levene’s test p<.05). For this reason, non-parametric comparisons for independent samples (Mann-Whitney U test) were used to compare differences across age groups. Analysis revealed that older adults employed greater head yaw rotation for pursuing the moving target compared to younger participants, but the effect was not so systematic (Mann-Whitney U = 175.0, p=.077, Fig 4c).

**Fig 4:**
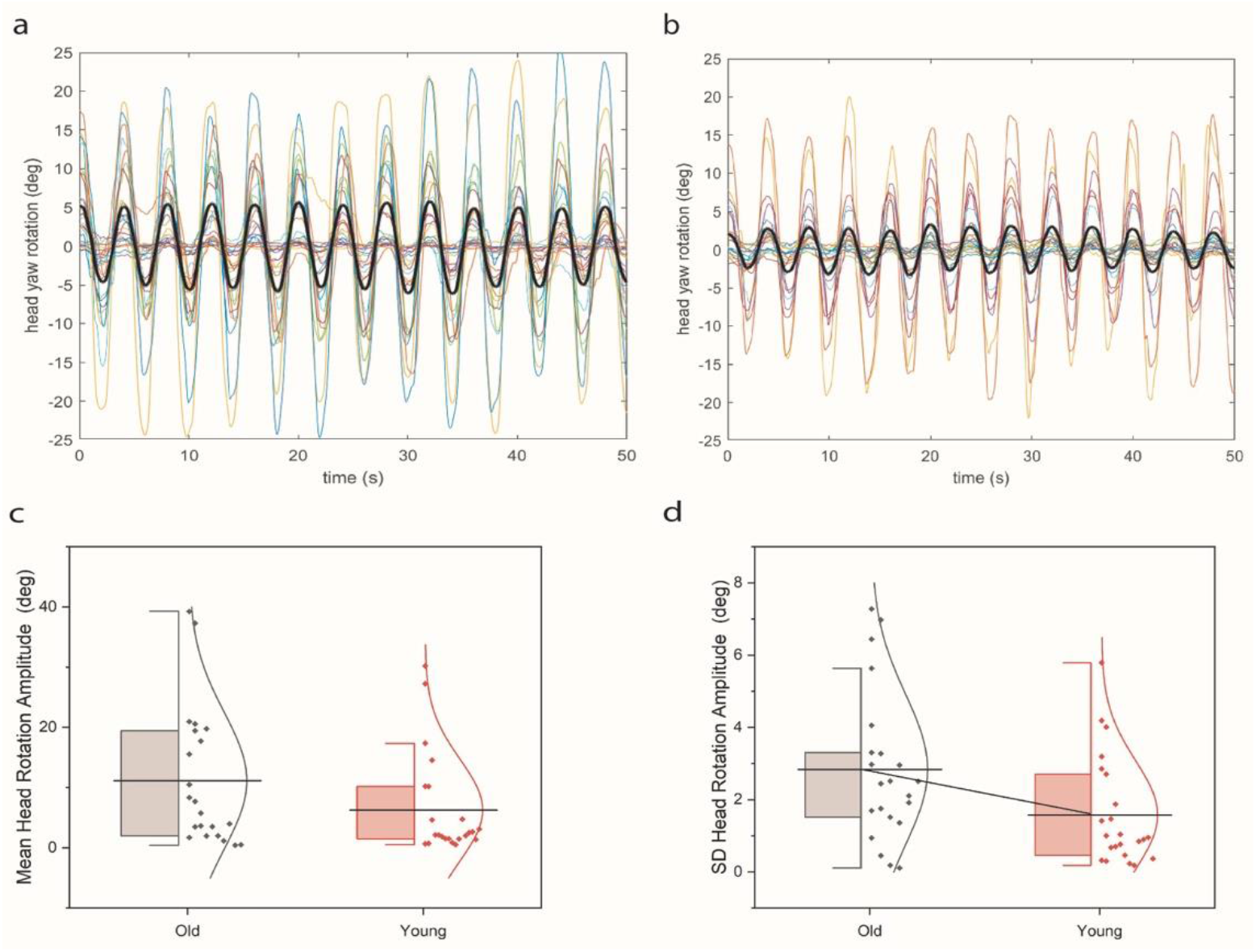
Individual time series of head yaw rotation plotted together with the group mean (thick black line) for the old (a) and the young (b) group during the gaze pursuit task. Mean (c) and standard deviation (d) of head yaw rotation across cycles *:< for young at p<.05

The SD of the head rotation amplitude across the target tracking cycles (Fig 4d) was normally distributed but only in the old group (Kolomogorov-Smirnov test: old group: p>.05, young group: p = .004). SD of head amplitude was also homogeneously distributed across groups (p>.05). Non-parametric between group comparison confirmed a significantly larger inter-cycle SD of head rotation amplitude for the old than the young group participants (Mann-Whitney U =152.0, p=.022).

The gaze-target peak CCF during the gaze pursuit task did not reveal a significant effect of age (Mann-Whitney U = 146.0 p>.05). All participants showed gaze-target peak CCF values greater than.8 (>80%).

### Relationship between head engagement in tracking and postural sway

To further investigate the possible relationship between head engagement during the gaze pursuit task and standing balance, we correlated the mean amplitude and SD of head yaw rotation with the postural measures during the gaze pursuit task. The scatterplots with the linear regression fit between the mean amplitude and SD of head yaw rotation and each of the postural sway variables is plotted in Fig 5 separately for the old and young group participants. These scatterplots revealed weak and non-significant relationships between the mean and SD of head rotation amplitude and sway measures with r values ranging from – 0.23 to 0.33 (p>.05).

**Fig 5:**
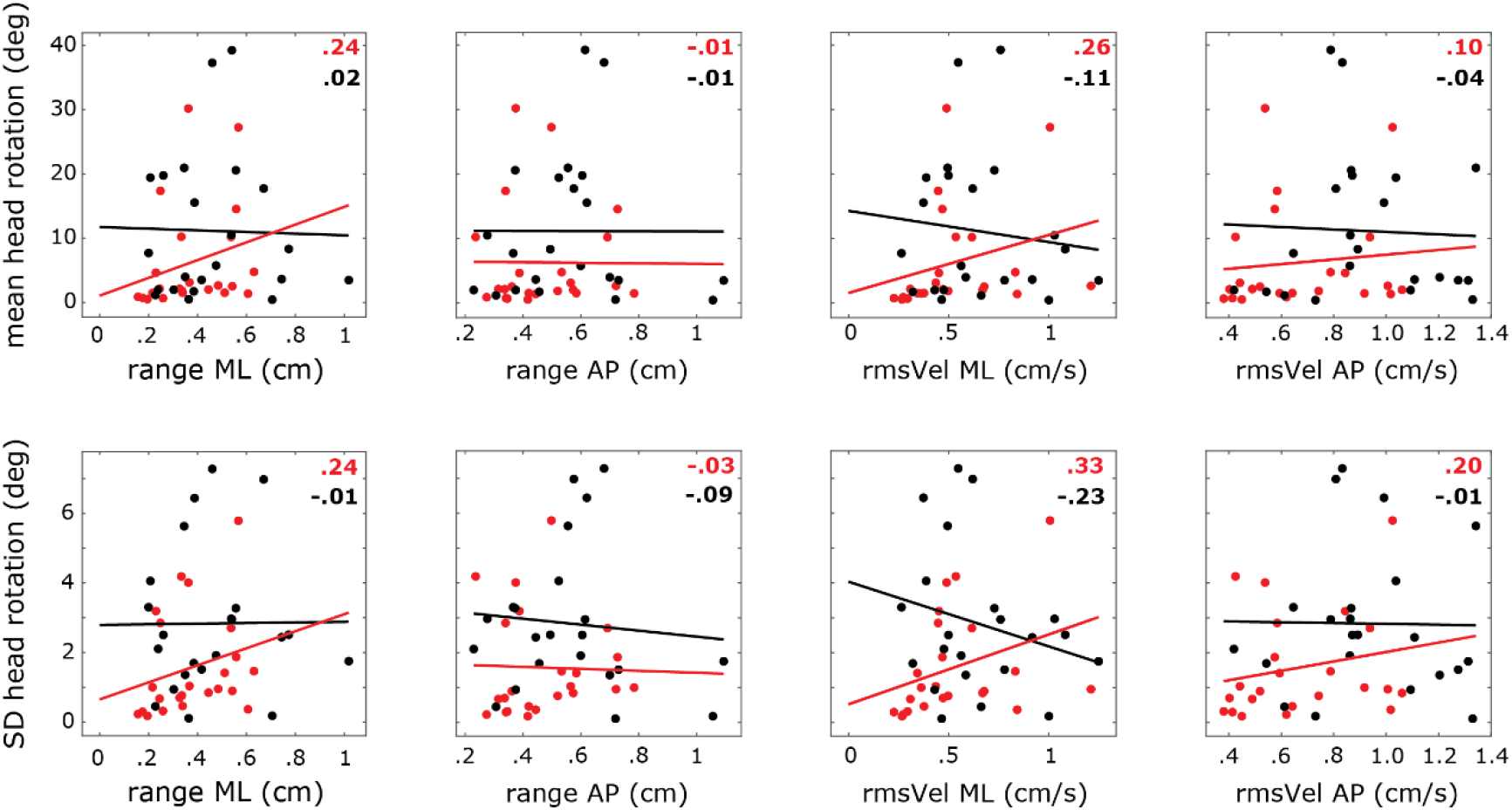
Scatterplots with linear regression lines between mean amplitude (a) and SD (b) of head yaw rotation and postural sway (CoP) variables for the young (red dots) and old (grey dots) group participants. Correlation coefficients (values on bottom of each sub-figure) were low and non-significant.

## Discussion

The results of the present study confirmed our first hypothesis suggesting that head free gaze pursuit of the horizontally moving target compromises standing balance to a greater extent in older than young adults relative to gaze fixation. However, despite the more variable head engagement of the older participants in the pursuit task, partially confirming our second hypothesis, there was no association between the amplitude and variance of head rotation and postural sway measures. Thus, the third hypothesis attributing the age-related greater sway to head yaw movements employed by older adults during the gaze pursuit task was not confirmed.

### The effect of the visual task on standing balance

Postural sway in the mediolateral (x) and anteroposterior (y) directions increased during the gaze pursuit compared to the fixation task. The present results confirm other work showing that smooth pursuit of a target moving on a static visual field destabilizes both static (Laurens et al. 2010; Thomas et al. 2016) and dynamic balance (Thomas et al. 2017) relative to gaze fixation. In addition, body sway is larger during free viewing than during gaze fixation (Thomas et al. 2016), also indicating that eye movements can compromise postural stability. This increased postural sway could be due to the retinal slip induced by the eye movement, for instance while following the moving target’s image. Preserving stability of a given visual input on the retina is important for the accurate detection of postural shifts (Schulmann et al. 1987). This however may necessitate more dynamic eye movements to stabilize the target’s image on the retina which could compromise stability (Glasauer et al. 2005). Extraretinal signals, such as eye proprioception and efference copy of the eye movement, are thought to improve balance by improving detection of body sway, but these are effective only for larger lateral body sway (for a review see Guerraz & Bronstein 2008).

Contrary to previous studies that restricted head motion during visual task performance by selecting a small target eccentricity (<15° of visual angle) (Stoffregen 2006), the present experimental paradigm imposed a larger visual angle of the target motion (≅ 31°) which required head engagement in the tracking task. This revealed an age group by task interaction on CoP excursion indicating that the destabilizing effect of gaze pursuit was greater in older than young participants, in contrast to the previously reported absence of age related differences on the postural sway increase between fixation and tracking (Thomas et al. 2016). It is not unreasonable to hypothesize that the greater, age-induced postural sway during the gaze pursuit task is due to larger head engagement in the target tracking task. In this case, inertial effects associated with the head’s movement may be affecting whole-body posture. In addition, extra flow of the visual field’s image due to the more variable head rotation (i.e., additional retinal slip) could have negative effects on standing balance. Yet, when rotating the head, additional sensory input from neck muscles and vestibular end organs can provide a more reliable estimate of gaze direction. Thus, moving the head could allow greater sensorimotor certainty and therefore improve body posture. To further explore these possibilities, we compared the mean amplitude and variance of head yaw rotation during the gaze pursuit task across the two age groups and investigated its possible impact on standing balance.

### Age related differences in head engagement and gaze strategy during visual tracking

Older adults pursued the horizontally moving target employing, descriptively, greater, and more variable head engagement than younger adults did, though only the later was statistically significant. Generally, older adults appear to engage their head during visuomotor task performance more than younger adults do. This has been found during vertical and horizontal saccades performed from a semi-tandem posture (Polastri et al. 2019) but also when re-orienting gaze during walking (Cinelli et al. 2008). This could be a compensation for age-induced limitations in positional precision (Kolarik et al. 2010; Maruta et al. 2017), range (Moschner & Baloh 1994; Paige 1994) and eye-target velocity gain during smooth pursuit tasks (Kanayama et al. 1994; Moschner & Baloh 1994; Matheron et al. 2008; Lee et al. 2019). Head rotation can allow for smaller and slower eye movements and decrease uncertainty about gaze orientation (Saglam et al. 2011) improving visuomotor accuracy. This idea is in line with the present results revealing a high cross-correlation between gaze and target motion (CCF > 0.8) regardless of age. Considering that the target was often followed also with a considerable head rotation, particularly by older adults, it could be argued that head rotation contributes to gaze orientation by providing extra sensory input, in line with previous postulations (Saglam et al. 2011). Previous work also confirms the absence of age-related deficits on visuo-motor accuracy when the head is free to move during gaze pursuit tasks performed in a natural environment (Dowiasch et al. 2015) or during visuo-postural coordination tasks that require tracking of periodically moving targets with the whole body (Sotirakis et al. 2016; Sotirakis et al. 2017). Yet, this high correlation may result from the fact that the target motion was temporally and spatially predictable, which allowed participants to anticipate the impeding target motion and predict the target’s spatial continuation based on previous experience, a skill that is preserved in old age (Chapman & Hollands 2007; Sprenger et al. 2011).

### The impact of head rotation on standing balance

We hypothesized that the greater head engagement is the source of the greater age-related sway observed during the tracking task compared to fixation. Specifically, we inferred that the increased head engagement might have led to inertial effects causing postural instability. Contrary to our expectation, analyses did not reveal an association between the head rotation variables and postural sway measures during the gaze pursuit task. Correlation coefficients were low, ranging from −0.23 to 0.33 and non-significant (Fig 5). The flat regression lines, for most of the sway measures, indicate that those, mostly older participants who engaged their head motion to track the moving target did not exhibit increased postural sway. Therefore, the extra head movement does not seem to be the reason for the increased sway noted in tracking compared to fixation and thus appears not to hamper balance control. This might be related to the fact that target motion was relatively slow (≅ 15°/s) compared to previous studies (> 20°/s; e.g., Paquette & Fung 2011b; Maruta et al. 2017), and so the induced head rotation during tracking in our study might have been relatively slow, minimizing inertial influences that could compromise postural stability. Alternatively, older adults may compensate for the extra head rotation by employing other control mechanisms such as ankle stiffening by increasing muscle co-contraction. An ankle stiffening strategy in anticipation of visual disturbances has previously been reported for older adults (Eikema et al. 2012). However, if a stiffening strategy was used to compensate for the extra head rotation, this should be reflected in a postural stabilization. Although the slight negative slope between the RMS of the CoP velocity and the two head rotation variables in the ML direction might point towards such a stabilization strategy, this idea is highly speculative because the overall net effect was to increase sway during tracking compared to fixation.

The degree of head engagement in visual tracking was characterized by large individual differences. A closer look at the individual head rotation curves plotted in Fig.5 reveals that several participants of the young group also pursued the moving target with considerable head rotation (>10° peak to peak amplitude). Therefore, head engagement during performance of precise visual tasks such gaze pursuit of a moving target might not be an age-specific strategy. Although further evidence is needed to explore this possibility, what can be confirmed by the non-significant correlations shown in Fig 5 is that also for young adult participants, head engagement in the visual tracking task does not explain the extra sway seen in tracking compared to fixation.

## Conclusions

Our study reveals, for the first time, that when the head is unrestrained during performance of visual tasks, gaze smooth pursuing of a horizontally moving target destabilizes standing balance to a greater extent in older than young individuals. We hypothesized that this may be due to an age-specific gaze strategy of engaging greater and more variable head yaw rotation in target pursuit possibly compensating for the age-related limited gain and velocity range of the eyes’ smooth pursuit movements. This hypothesis was not confirmed by the present results which did not demonstrate an association between head rotation amplitude and variance during target tracking and postural sway measures in older adults. This suggests that posture is robust against the more variable head rotation engaged by older adults when tracking a moving target. Yet, this leaves open the question why head free gaze pursuit of a horizontally moving target compromises standing balance to a greater extent in older than young adults compared to gaze fixation. Future investigations should focus on the effect of gaze pursuit under more challenging balance conditions in older adults and provide more extensive understanding on the control of dynamic balance while performing complex visual skills. This could result in extracting useful suggestions to improve exercise means and techniques used in fall prevention.

## Acknowledgments

We thank Dr. Evagelia Kouidi for her advice and help in older adults’ recruitment and health screening, as well as Iason Christodoulou and Andriana Teloudi for their valuable assistance in data collection. DV is supported by the German Research Foundation (Deutsche Forschungsgemeinschaft) under grant agreement VO 2542/1.

